# Feed efficiency of lactating Holstein cows was not as repeatable across diets as within diet over subsequent lactation stages

**DOI:** 10.1101/2021.02.10.430560

**Authors:** Amélie Fischer, Philippe Gasnier, Philippe Faverdin

**Author notes:** Corresponding author: Amélie Fischer –.

## Abstract

**Background:** Improving feed efficiency has become a common target for dairy farmers to meet the requirement of producing more milk with fewer resources. To improve feed efficiency, a prerequisite is to ensure that the cows identified as most or least efficient will remain as such, independently of diet composition. Therefore, the current research analysed the ability of lactating dairy cows to maintain their feed efficiency while changing the energy density of the diet by changing its concentration in starch and fibre. A total of 60 lactating Holstein cows, including 33 primiparous cows, were first fed a high starch diet-low fibre (diet S^+^F^-^), then switched over to a low starch diet-high fibre (diet S^-^F^+^). To know if diet affect feed efficiency, we compared the ability of feed efficiency to be maintained within a diet over subsequent lactation stages, known as repeatability of feed efficiency, with its ability to be maintained across diets, known as reproducibility of feed efficiency. To do so we used two indicators: the estimation of the error of repeatability/reproducibility, which is commonly used in metrology, and the coefficient of correlation of concordance (CCC), which is used in biology. The effect of diet change could also lead to a change in cows sorting behaviour which could potentially affect feed efficiency if for example the most efficient cows select more concentrate than the least efficient. We therefore analysed the relationship between the differences in individual feed refusals composition and the differences in feed efficiency. To do so, the composition of each feed refusal was described with its near infra-red (NIR) spectroscopy and was performed on each individual feed ingredient, diet and refusals and used as composition variable. The variability of the NIR spectra of the refusals was described with its principal components thanks to a principal component analysis (PCA). The Pearson correlation was estimated to check the relationship between feed efficiency and refusals composition, i.e. sorting behaviour.

**Results:** The error of reproducibility of feed efficiency across diets was 2.95 MJ/d. This error was significantly larger than the errors of repeatability estimated within diet, which were 2.01 MJ/d within diet S^-^F^+^ and 2.40 MJ/d within diet S^+^F^-^. The CCC was 0.64 between feed efficiency estimated within diet S^+^F^-^ and feed efficiency estimated within diet S^-^F^+^. This CCC was smaller than the one observed for feed efficiency estimated within diet between two subsequent lactation stages (CCC = 0.72 within diet S^+^F^-^ and 0.85 within diet S^-^F^+^). Feed efficiency was poorly correlated to the first two principal components, which explained 90% of the total variability of the NIR spectra of the individual refusals. This suggests that feed sorting behaviour did not explain differences in feed efficiency.

**Conclusions:** Feed efficiency was significantly less reproducible across diets than repeatable within the same diet over subsequent lactation stages, but cow’s ranking for feed efficiency was not significantly affected by diet change. This loss in repeatability across diets could be due to a more pronounced feed sorting subsequent to the change in diet composition. However, the differences in sorting behaviour between cows were not associated to feed efficiency differences in this trial neither with the S^+^F^-^ diet nor with the S^-^F^+^ diet. Those results have to be confirmed on diets having different forage to concentrate ratios to ensure that the least and most efficient cows will not change.

## BACKGROUND

To be more competitive, dairy farmers have to increase the efficiency of resources use while reducing their environmental footprint. With an expected increase of world population (United Nations, 2017), feeds for dairy cow may shift towards more non-human edible feeds. The challenge for dairy farmers will be to improve feed efficiency, while facing more volatile feed prices (HLPE, 2011) and lower availability of feeds, which are directly edible by human. Dairy cows’ diets will therefore become more variable in the future. Feed efficiency can be improved either by selecting the most efficient cows, thanks to an index including feed efficiency, or by improving feed efficiency of the least efficient cows with precision feeding. If feed efficiency is included as a genomic selection trait, feed efficiency has to be reproducible independently of diet and environment. The sensitivity of dairy cow’s ranking for feed efficiency to diet composition, also called interaction between genetic and environment (Hill and Mackay, 2004), needs to be evaluated to know if some cows perform better and some worse when changing diet composition.

Repeatability is defined as the capacity of a method to give the same results when using the same sample and repeating measurements in the same experimental conditions (JCGM, 2012), that is within diet when applied to efficiency. Reproducibility is a repeatability done while changing one specific characteristic in the experimental conditions (temperature, diet, operator; (JCGM, 2012)), that is for example by changing diet’s composition. When estimating reproducibility of an indicator under different environmental conditions, it is therefore essential to compare it with its repeatability under similar conditions to isolate errors associated with diet change from errors associated with the method. In literature, the reproducibility of feed efficiency when changing the diet was lower compared to the repeatability within a given diet, as estimated with the correlations of residual feed intake (RFI) within and across diets (r = 0.33 vs 0.42 in steers, (Durunna et al., 2011); r = 0.44-0.64 vs 0.53-0.70 in dairy cows, (Potts et al., 2015)). Animals will thus not necessarily rank the same for feed efficiency when changing diet composition. Repeatability estimation, as defined by the proportion of genetic and permanent environment variances within and across lactations in the total variance of RFI, is highly variable across studies and countries with values between 0.47 and 0.90 (Connor et al., 2013; Tempelman et al., 2015). This variability across studies is associated with differences in diet composition and in period length between studies. Indeed, in lactating dairy cows, the correlation of short-term feed efficiency with full lactation feed efficiency increases with later lactation stages and longer periods (Connor et al., 2019). The comparison between repeatability and reproducibility needs therefore feed efficiency to be estimated over a long enough period to get a robust estimation within diet. Feed efficiency is generally estimated with diets offered ad libitum with a minimum amount of refusals (in general 5 – 10% of offered). Cows can therefore potentially perform feed sorting, which may lead to a consumed diet that differs from the distributed diet both in composition and in nutritive value. This difference between refusals composition and distributed diet composition has therefore to be considered when analysing the change in feed efficiency while changing diet’s composition because feed sorting behaviour could affect feed efficiency. For instance, Dykier et al. (2020) observed that RFI was negatively correlated to the intake of long particles (r = −0.30, p < 0.05) and positively correlated to intake of short particles (r =0.22, p < 0.1) in beef steers fed ad libitum. This difference in sorting behaviour resulted in a consumed diet which concentration in crude protein increased with RFI (r = 0.25, p < 0.05). Differences in feed composition were characterized by particle size differences in Dykier et al. (2020). However the method for particle size composition (Lammers et al., 1996; Kononoff et al., 2003) is burdensome and time consuming. The advent of near-infrared (NIR) spectroscopy opens new ways to determine diet or feed compositions at high throughput.

Indeed the NIR spectrum is sensitive to physical and chemical characteristics of the sample, and has therefore been used to determine nutritive value of feed, but also to discriminate samples according to their composition (Pérez-Marín et al., 2004; De la Roza-Delagado et al., 2007; Li et al., 2007).

The main objective of the current study was therefore to check the ability of feed efficiency to be maintained across different diets. To achieve those objectives a trial was set up with lactating dairy cows that were fed with two diets. These diets differed in energy density, by lowering the starch concentration and increasing the fibre concentration of the diet. The feed efficiency was estimated within diet using the method developed in a previous paper (Fischer et al., 2018). The novelty of the current paper is to estimate feed efficiency reproducibility across diets by combining two methods: the commonly used CCC in biology and the comparison of the error of reproducibility across diets with the error of repeatability within diet, as commonly used in metrology (JCGM, 2012). Indeed, to estimate if FE is maintained across diets, its reproducibility across diets has to be compared to its repeatability within diet. If the reproducibility results are as good as the repeatability results within diet, then one can conclude that FE is as repeatable across diets as it is within diet. Opposedly if the reproducibility results are worse than repeatability within diet, then one can conclude that the ability of FE to be maintained across diets is not as good as within diet. As highlighted in the previous paragraph, a diet change could also lead to a change in cows sorting behaviour which could potentially affect feed efficiency. A second objective was therefore to check that the change in feed efficiency associated with diet change was not explained by differences in sorting behaviour. We therefore checked that feed efficiency was not associated with feed sorting behaviour by analysing the feed composition of each cow’s diet refusals with NIR spectroscopy.

## MATERIAL & METHODS

### Experimental Design

The experimentation was performed at the INRAE-Institut Agro UMR PEGASE research facility of Mejusseaume (Le Rheu, France). An initial group of 68 Holstein cows were housed in a free-stall barn with free access to water. These cows were monitored for feed efficiency from calving (grouped between end of August and middle of October 2017) to end of June 2018, near the end of the lactation. A sequential design was adopted to feed the cows with a first diet that was high in starch and low in Fibre (diet **S^+^F^-^**) and a second diet that was low in starch and high in fibre (diet **S^-^F^+^**). The protein content was adjusted in diet S^-^F^+^ to have a ratio between energy and protein that was similar to the ratio in diet S^+^F^-^. The diet S^+^F^-^ was fed from calving to March 18^th^ 2018, then all cows switched over to diet S^-^F^+^ until end of June 2018. To make the feed efficiency estimated over the two feeding periods comparable, the same period length was chosen for both periods. Each diet was fed for at least 95 days.

The first 23 days subsequent to the change in diet from S^+^F^-^ to diet S^-^F^+^ were considered as an adaptation period to the new diet and were therefore removed from the dataset. Each period included the last 72 days of data. 62 cows had data over both 72-day periods and were therefore kept for further analysis. Each period has been split in two sub-periods of 36 days (Figure 1) to be able to estimate a repeatability within diet and to reach a supposedly correlation with full lactation RFI of at least 0.8 according to Connor et al. (2019).

**Figure 1.**
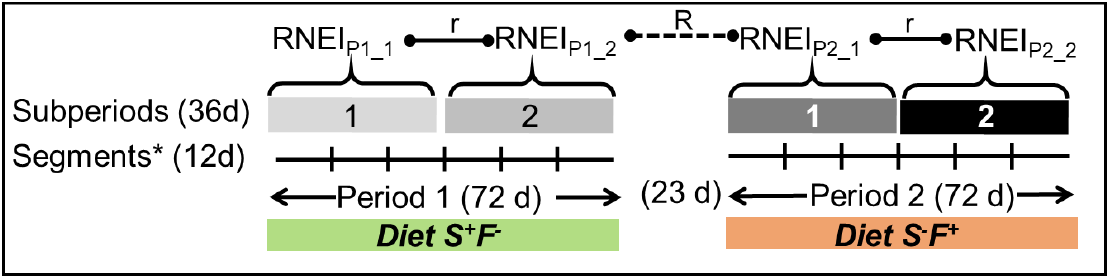
Diagram of the design used to characterize the repeatability (r) and reproducibility (R) of residual net energy intake (RNEI). The repeatability compares RNEI within diet; it is shown with the solid line 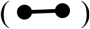. The reproducibility compares RNEI across diets; it is shown with the dashed line 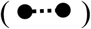. The dairy cows started first with the diet high in starch and low in fibre (S^+^F^-^) and switched over to the diet low in starch and high in fibre (S^-^F^+^).

Reproducibility was estimated using the last 36-days of the first period and the first 36-days of the second period to be comparable and compared with repeatability estimation over the two 36-days segments within diet (Figure 1).

### Phenotypic Measurements

#### Individual feed intake and feed nutrient analysis

Cows were fed individually twice a day after each milking (7:00 and 16:00). Daily intake was estimated individually as the difference between daily offered feed and next morning’s refusals. The diet was offered ad libitum to maintain an average of 10% refusals per cow. Each cow was fed in her own manger, only accessible by one cow thanks to the identification chip on her neck collar. Each feed has been sampled once a week for concentrates, and once a day for the forages to estimate dry matter, which was used to estimate individual feed dry matter intake (DMI). A bulk sample was taken for each ingredient and silo for nutrient value analysis. Diet S^+^F^-^ included maize silage, soybean meal, dehydrated alfalfa and a mix of energy concentrates, minerals and vitamins (Table 1). Diet S^-^F^+^ was formulated to have a lower starch concentration by replacing starch with fibre while keeping a similar ratio between metabolizable protein and net energy for lactation than in diet S^+^F^-^. To do so, wheat straw was added to the ingredients included in diet S^+^F^-^ to replace the energy concentrate and part of soybean meal and maize silage (Table 1). In addition to both diets, cows had access to a gas emissions monitoring system, the Greenfeed^®^ (see “Individual performance: milk, body weight and body condition, methane and carbon dioxide emissions” section), which distributes small drops of energy concentrates to maintain the cow in the gas recording system. The amount of energy concentrates distributed per cow per day in the Greenfeed^®^ station was added to the daily intake at the manger. All feed samples were freeze-dried and ground with a 3-blade knife mill through a 0.8-mm screen. The organic matter content was determined by ashing for 5h at 550°C in a muffle Furnace (Association Française de Normalisation, 1997). The concentrations of neutral and acid detergent fibre were measured according to Van Soest et al. (1991) using a Fibersac extraction unit (Ankon Technology, Fairport, NY, USA). Fat content was measured by ether extraction and starch analyses were performed by polarimetry (LABOCEA, Ploufragan, France). Nitrogen concentration for all samples was determined by the Dumas method (Association Française de Normalisation, 1997) with a LECO Nitrogen Determinator (Leco, St Joseph, MI, USA). Nutritive values of feeds were calculated from their chemical composition according to equations in INRA (2010). Diet S^+^F^-^ had 226 g starch/kg DM, 351 g neutral detergent fibre/kg DM for a net energy concentration of 6.62 MJ/kg DM and 105 g metabolizable protein/kg DM. Diet S^-^F^+^ was 19% lower in starch concentration and 16.4% higher in neutral detergent fibre concentration than diet S^+^F^-^ for a net energy concentration of 5.98 MJ/kg DM and 93 g metabolizable protein/kg DM (Table 1).

**Table 1.** Composition of both experimental diets (S^+^F^-^ and S^-^F^+^) and their chemical analysis.

|  | Diet S<sup>+</sup>F<sup>-</sup> | Diet S<sup>-</sup>F<sup>+</sup> |
| --- | --- | --- |
| Feed composition (% DM +/- SD<sup>1</sup>) |  |  |
| Maize silage | 65.0 +/- 1.6 | 59.1 +/- 0.9 |
| Soybean meal | 17.8 +/- 0.9 | 13.3 +/- 0.3 |
| Dehydrated lucerne | 8.1 +/- 1.6 | 14.4 +/- 0.2 |
| Energy concentrate<sup>2</sup> + Minerals and vitamin complement<sup>3</sup> | 9.1 +/- 1 | 1.6 +/- 0.7 |
| Wheat straw | 0 | 11.6 +/- 0.3 |
| Diet analysis (+/- SD<sup>1</sup>) |  |  |
| DM (%) | 42.9 +/- 0.7 | 47.1 +/- 0.3 |
| OM (%) | 94.0 +/- 0.15 | 94.1 +/- 0.04 |
| Crude protein (g/kg) | 167 +/- 4.9 | 145 +/- 1.1 |
| ADF (g/kg) | 198 +/- 2.9 | 249 +/- 1.2 |
| NDF (g/kg) | 351 +/- 3.6 | 420 +/- 1.6 |
| Starch (g/kg) | 226 +/- 3 | 183 +/- 2.2 |
| Net energy for milk (MJ/kg) | 6.62 +/- 0.107 | 5.98 +/- 0.014 |
| Metabolizable protein (g/kg) | 105 +/- 1.8 | 93 +/- 0.4 |

<sup>1</sup>SD were calculated using the day-to-day change in offered diet composition on an individual cow
basis. <sup>2</sup>The part of energy concentrate in the diet, as described here, includes the part of energy
concentrates distributed at the Greenfeed® station. <sup>3</sup>Concentrates included 17.8% wheat, 17.8%
maize, 17.8% sugar beet pulp, 17.8% barley, 13% wheat bran, 3% beet molasses, 0.9% oil, 0.9% salt,
11% minerals and vitamin complement (including 6% phosphorus, 24% calcium, 5% magnesium, and
other minerals and vitamins).
DM = dry matter; OM = organic matter; ADF = acid detergent fibre; NDF = neutral detergent fibre

#### Individual performance: milk, body weight and body condition, methane and carbon dioxide emissions

Milk yield was recorded at each milking with milk meters (DeLaval, Tumba, Sweden). Milk fat and protein concentrations were analysed by mid infrared spectrometers (Lillab, Chateaugiron, France) from morning and afternoon milk samples of two days per week. Milk fat and milk protein concentrations were calculated as weighted averages relatively to the morning and afternoon milk production of the day of sampling. Cows were weighed automatically after each milking (W-2000, DeLaval, Tumba, Sweden) to get an empty udder body weight (**BW**). All cows were scored for body condition once a month by 3 trained persons according to the scale developed by Bazin (1984), going from 0 for an emaciated cow to 5 for a fat cow with 0.25 unit increments.

Methane emissions were recorded with two Greenfeed^®^ units (C-Lock Inc., Rapid City, SD, USA). A Greenfeed^®^ unit is designed as a dispenser of concentrates to measure methane and carbon dioxide emissions each time a cow visits the feeder, therefore both methane and carbon dioxide were also monitored. A maximum of 720 g (30g/drop) concentrates was offered daily at the Greenfeed^®^ to attract cows in the Greenfeed. Each Greenfeed^®^ unit records methane emissions for up to 23 cows. Given the barn configuration and the two Greenfeed^®^ units, only 42 cows could therefore be monitored for methane emissions during the study. On average the cows visited the Greenfeed 2.2 /d/cow (+/− 0.9). Energy concentrates distributed at the Greenfeed^®^ units were included in the estimation of individual daily feed intake.

### Checking for feed sorting behaviour

Individual refusals were sampled by collecting about 0.5 to 1 kg of fresh weight refusals per cow once a week for 6 weeks during each of the two experimental periods to check if the change in diet’s feed ingredients between diet S^+^F^-^ and diet S^-^F^+^ modified cow’s feed sorting behaviour. The samples were dried in a forced-air oven at 60°C for 48 h. The refusals samples of each cow were pooled within period to end up with one sample per cow and per period, and ground through a 3-blade mill (0.8 mm screen). The same process was applied for each feed ingredient and each diet to estimate the difference in composition between feeds and refusals. Instead of determining each sample’s feed composition, we used an indirect approach based on near-infra red (NIR) spectroscopy. Indeed, we have seen in the background section of this paper that NIR spectroscopy can be used to differentiate samples differing in ingredients proportion. By definition, if the samples differ on a physical or chemical basis, their spectra will also be different. In this study, the differences in refusals composition and diets composition will be analysed through their NIR spectra, without estimating or analysing their chemical or physical characteristics. A Fourier transform near-infrared (FT-NIR) spectroscopy with MPA (Bruker Optik Gmbh, Ettligen, Germany) was used to characterize the spectra (from 3595 to 12490 cm^-1^, resolution 16 cm^-1^) of each individual feed, diet and refusal sample. A principal component analysis (PCA) was fitted on the spectra of the feeds, both diets and all refusals to summarize the dataset into principal components. A second PCA was performed on the spectra of the refusals only, with randomRNEI as a supplementary variable, to summarize the spectra of the refusals into fewer variables. The relationship between refusals composition and feed efficiency was estimated with the coefficient of correlation between randomRNEI and each of the principal component of this second PCA. Both PCA were performed with the FactomineR (Lê et al., 2008) and Factoshiny (Vaissie et al., 2020) packages of R (R Core Team, 2018).

### Outlier Detection

Among the 62 cows, 2 cows had issues with their manger and were therefore removed from the dataset because their intake data were not reliable enough, to end up with a group of 60 cows, including 33 primiparous cows. For methane and carbon dioxide data, a least rectangles regression of carbon dioxide emission against methane emission was fitted to detect methane outliers using the least.rect function of package RVAideMemoire (Hervé, 2018) in R (R Core Team, 2018). The data outside the range of three standard deviations of the residuals around the regression line were considered as outliers, and were removed. On average 0.9% (SD = 1.3%) of the initial methane data were removed per cow for being outliers. Methane data were then averaged per experimental period and per cow.

### Variables Calculation to Estimate Feed Efficiency

Estimation of feed efficiency requires DMI data, as well as all energy outputs or energy inputs to be considered for a lactating dairy cow. Energy outputs gather net energy in milk, energy required for maintenance, energy gained as adipose tissue and energy required for gestation. Other energy inputs include adipose tissue mobilization.

Net energy in milk was calculated according to the following equation (Faverdin et al., 2010):

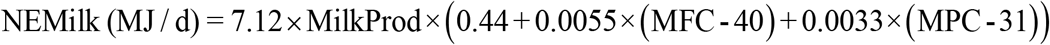

Where MilkProd is the milk production in kg/d, MFC is the milk fat concentration in g/kg and MPC is the milk protein concentration in g/kg.

Gestation requirement were estimated with the following equation defined by Faverdin et al. (Faverdin et al., 2010):

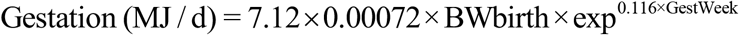

Where BWbirth is calf’s weight at birth and assumed to be 40 kg, GestWeek is the week of gestation.

Morning BW data used were smoothed with a moving average of the 15% neighbouring data, to better reflect change in maintenance and to be less sensitive to daily gutfill change. Monthly body condition score (**BCS**) data were filled to get daily BCS using a cubic Spline with the function smooth.spline in R (R Core Team, 2018) using each scoring day as a knot. Maintenance requirements were estimated with the metabolic BW, using the smoothed BW data, and calculated as BW^0.75^. Energy gained and energy mobilized as body reserves were estimated as the day-to-day change in smoothed BW. If the change was positive, it was attributed to body reserves gain, and body reserves loss was null. Conversely, if the change was negative, it was attributed to body reserves mobilization, and body reserves gain was null. Both BW gain and BW loss were constructed to be positive variables. Both, BW gain and BW loss, were multiplied by daily BCS to account for body reserves differences within a given BW change, resulting in the variables **BWlossBCS** and **BWgainBCS**.

### Estimation of feed efficiency

Feed efficiency was estimated as the residual feed intake with the method developed in a previous paper (Fischer et al., 2018). Briefly, instead of being estimated as the residual of the multiple linear regression that estimates the observed DMI with the main energy outputs and inputs, RFI was defined with a mixed model as the repeatable animal effect. Applied to our study, each sub-period of 36 days was subdivided in segments of 12 days to end up with three repeated measures for each cow within each sub-period (Figure 1). The initial model explained net energy intake (NEI) with the fixed effects of net energy in milk, metabolic BW, BWlossBCS, BWgainBCS, gestation requirement, BCS, their interaction with parity and sub-period, and the fixed effect of parity, sub-period, and 12-day segment nested within sub-period. This model included the repeated effect of cow across the 12-day segments within sub-period and the random effect of cow within sub-period, and were grouped within sub-period. Only significant (p ≤ 0.05) interactions and variables were kept in the model. As in our previous paper (Fischer et al., 2018), feed efficiency was defined as the random part of the intercept of the mixed model 1 below, that was performed using PROC MIXED in SAS (Version 9.4 of the SAS System for Linux. 2017. SAS Institute Inc., Cary, NC, USA) with a heterogeneous autoregressive variance covariance matrix for the repeated statement. The variables were averaged per 12-day segment.

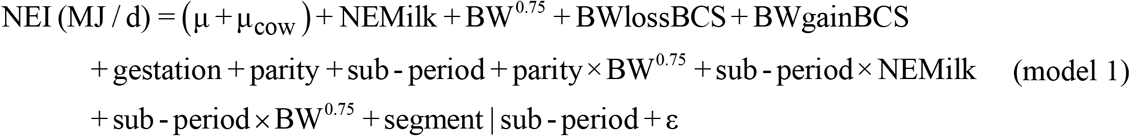

where NEMilk is the net energy in milk in MJ/d, BW^0.75^ is the metabolic BW in kg^0.75^, gestation is the gestation requirement in MJ/d, parity is the fixed effect of parity, sub-period is the fixed effect of sub-period, segment|sub-period is the fixed effect of 12-day segment nested within sub-period, parity×BW^0.75^ is the interaction between parity and BW^0.75^, sub- period×NEMilk and sub-period×BW^0.75^ are the interactions between sub-period and NEMilk, and sub-period and BW^0.75^, μ is the fixed intercept and μ_cow_ is the random part of the intercept and ε is the error. Feed efficiency was defined as μ_cow_ in model 1 and will be called random residual net energy intake **(RandomRNEI)**.

### Repeatability and Reproducibility of Feed Efficiency

Repeatability and reproducibility were estimated with 2 methods: the standard deviation of repeatability and standard deviation of reproducibility as defined by ICAR (JCGM, 2012), and Lin’s coefficient of correlation of concordance (CCC) (Lin, 1989).

#### Estimating Error of Repeatability and Reproducibility

Repeatability was estimated within diet with an analysis of variance. For repeatability the model 3 below of analysis of variance was fitted once with the data of feed efficiency within diet S^+^F^-^ to get the repeatability within diet S^+^F^-^, and once with the data of diet S^-^F^+^ to get the repeatability within diet S^-^F^+^. Reproducibility was estimated with an analysis of variance with the data of the second sub-period when cows were fed diet S^+^F^-^ and the data of the first sub-period when cows were fed the diet S^-^F^+^, to be able to estimate the variance associated with diet change. Both analysis of variance were performed using the Anova function of car package (Fox and Weisberg, 2011) in R (R Core Team, 2018) as follows:

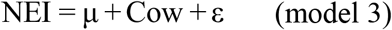

Where Cow stands for the fixed effect of cow, μ is the intercept and ε is the error. Standard deviations of repeatability and of reproducibility were defined as the standard deviation of ε in model 3. An F-test was performed with the var.test function in R (R Core Team, 2018) to test if the error of repeatability in diet S^+^F^-^ was similar to the error of reproducibility across diets, and similarly by comparing error of repeatability in diet S^-^F^+^ with the error of reproducibility across diets.

#### Estimation of Repeatability and Reproducibility Correlations

The Lin’s coefficient of correlation of concordance was estimated between the feed efficiency estimated at two different sub-periods within diet for repeatability, and between feed efficiency estimated during the second sub-period in diet S^+^F^-^ and first sub-period in diet S^-^F^+^ for reproducibility. The CCC were estimated with the epi.ccc function of epiR package (Stevenson et al., 2020) in R (R Core Team, 2018).

All statistical analysis were done with the significance level of 0.05 (p ≤ 0.05).

## RESULTS

### Period Effect on Cows Performance

The decrease in dietary net energy for lactation and in metabolizable protein was confounded with the increase of lactation stage as the experimentation was based on a sequential design. Therefore when effect of diet is mentioned here, it is confounded with the effect of lactation stage.

The diet change decreased net energy intake, without significantly changing dry matter intake (p = 0.26). Indeed, Cows ate on average 23.4 kg DM/d over both periods. They ate 156 MJ/d when they were fed with diet S^+^F^-^, which was 13.9 % (19 MJ/d) more net energy than when they were fed with diet S^-^F^+^ (p < 0.01; Table 2).

**Table 2.** Dry matter intake, performance, methane emissions and efficiency of the 60 Holstein cows fed S^+^F^-^ diet first and then S^-^F^+^ diet three months later.

|  | Least squares means |  | Residual SD | p-value |
| --- | --- | --- | --- | --- |
|  | Diet S<sup>+</sup>F<sup>-</sup> | Diet S<sup>-</sup>F<sup>+</sup> |  |  |
| n (primiparous) | 60 (33) | 60 (33) |  |  |
| Intake |  |  |  |  |
| DMI (kg/d) | 23.7 | 23.1 | 2.94 | 0.26 |
| NEI (MJ/d) | 156 | 137 | 18.58 | <0.01 |
| Performance |  |  |  |  |
| Milk production (kg/d) | 36.4 | 29.9 | 4.93 | <0.01 |
| Milk Fat production (g/d) | 1,387 | 1,154 | 194.0 | <0.01 |
| Milk Protein production (g/d) | 1133 | 933 | 145.6 | <0.01 |
| Milk Protein concentration (g/kg) | 31.4 | 31.4 | 2.44 | 0.88 |
| Milk Fat concentration (g/kg) | 38.5 | 39.0 | 4.27 | 0.56 |
| NE in Milk (MJ/d) | 111.4 | 92.1 | 14.17 | <0.01 |
| BW (kg/d) | 651 | 679 | 68.2 | 0.03 |
| BCS | 1.96 | 2.08 | 0.23 | <0.01 |
| BW Gain (kg/d) | 0.40 | 0.37 | 0.16 | 0.28 |
| BW Loss (kg/d) | 0.14 | 0.32 | 0.12 | <0.01 |
| Methane |  |  |  |  |
| Methane production (g/d) | 510 | 523 | 66.2 | 0.41 |
| Methane yield (g/kg DMI) | 21.4 | 22.6 | 2.39 | 0.03 |
| Methane yield (g/kg Milk) | 14.0 | 17.6 | 2.58 | <0.01 |

DMI: dry matter intake; NEI: net energy intake estimated using equation in Faverdin et al. [12]; NE in milk: net energy in milk estimated using equation in Faverdin et al. [12]; BW: body weight; BCS: body condition score; CH<sub>4</sub>: methane emission. Diet S<sup>+</sup>F<sup>-</sup>: high in starch and low in fibre; diet S<sup>-</sup>F<sup>+</sup>: low in starch and high in fibre.

Cows partitioned 92.1 MJ/d in milk on diet S^-^F^+^, which was 17.3% (19.3 MJ/d) less than with diet S^+^F^-^ (p < 0.01, Table 2). This difference in net energy exported in milk between diets was also observed for milk production. Indeed, cows produced 29.9 kg milk/d with diet S^-^F^+^, which was 17.9% (6.5 kg/d) less than with diet S^+^F^-^ (p < 0.01, Table 2). Change in dietary net energy and metabolizable protein concentrations did neither significantly affect milk protein concentration (p = 0.88) nor milk fat concentration (p = 0.56), with average concentrations of 31.4 g protein/kg milk and 38.8 g fat/kg milk (Table 2). Given the steady milk content and a decreasing milk production, the decrease in net energy and metabolizable protein in the diet decreased milk protein production (p < 0.01) and milk fat production (p < 0.01).

Maintenance related variables, known as BW and BCS, were lower when cows were fed diet S^+^F^-^ (p = 0.03 for BW and p < 0.01 for BCS), which was also at earlier lactation stages, with averages of 651 kg and 1.96 BCS with diet S^+^F^-^, and 680 kg and 2.08 BCS with diet S^-^F^+^ (Table 2).

Variables associated with body reserves change, identified as BW loss and BW gain in table 2, were differently affected by diet. Cows mobilized more BW when they were fed with S^-^F^+^ diet, also known as period 2, with a loss of 0.32 kg/d, than when fed with S^+^F^-^ diet, also known as period 1, with a loss of 0.14 kg/d (p < 0.01, Table 2). Gain in BW did not significantly differ between both diets, with an average gain of 0.39 kg/d.

Dietary decrease in starch, replaced with fibre, confounded with the effect of lactation stage, did not significantly change methane emissions, as cows emitted on average 517 g methane/d (Table 2). Nevertheless, this dietary change increased methane yield from 21.4 g methane/kg DMI up to 22.6 g methane/kg DMI (p = 0.03) and from 14.0 g methane/kg milk up to 17.6 g methane/kg milk when switching from diet S^+^F^-^ to diet S^-^F^+^ (p < 0.01, Table 2).

### Effect of Diet change on Feed Efficiency

Feed efficiency was more variable when cows were fed the S^-^F^+^ diet than when fed the S^+^F^-^ diet. Feed efficiency, as estimated with randomRNEI, had a standard deviation of 4.49 MJ/d in sub-period 1 and 4.61 MJ/d in sub-period 2 when cows were fed S^+^F^-^ diet, and 5.18 MJ/d in sub-period 1 and 5.21 MJ/d in sub-period 2 when cows were fed the S^-^F^+^ diet. The change in randomRNEI induced by diet change (randomRNEI diet S^+^F^-^ - randomRNEI diet S^-^F^+^) was negatively correlated with the change in methane yield, as per kg DMI, (CH4/DMI diet S^+^F^-^ - CH4/DMI diet S^-^F^+^) with a Pearson correlation of - 0.31 (p = 0.05), but was neither significantly correlated with the change in methane production per day (p = 0.12) nor with the change in methane yield, as per kg milk (p = 0.98). This means that a cow that had a lower randomRNEI (higher feed efficiency) in diet S^-^F^+^, also had a higher methane yield per kg DMI in diet S^-^F^+^ than when fed with the S^+^F^-^ diet, and conversely.

### Feed Efficiency Reproducibility across Diets

Cows’ feed efficiency was not as reproducible across diets than repeatable within diet (Table 3 and Figure 2). Errors of reproducibility, when comparing efficiency across diets, were larger than the errors of repeatability within diet, regardless of diet (Table 3). Indeed, the reproducibility error of randomRNEI that was 2.95 MJ/d was significantly larger (p < 0.05, Table 3) than the repeatability errors for diet S^+^F^-^ and for diet S^-^F^+^ that were 2.40 and 2.01 MJ/d, respectively (Table 3). This lower reproducibility across diets compared with repeatability within diet tended to be observed with the CCC when comparing reproducibility across diets with repeatability in diet S^-^F^+^, but not when compared with repeatability within diet S^+^F^-^ (Table 3). Indeed, the CCC between randomRNEI estimated in diet S^+^F^-^ and randomRNEI estimated in diet S^-^F^+^ was 0.64, which was smaller compared with the correlations of 0.72 (p = 0.55) within diet S^+^F^-^ and 0.85 (p = 0.055) within diet S^-^F^+^ (Table 3).

**Figure 2.**
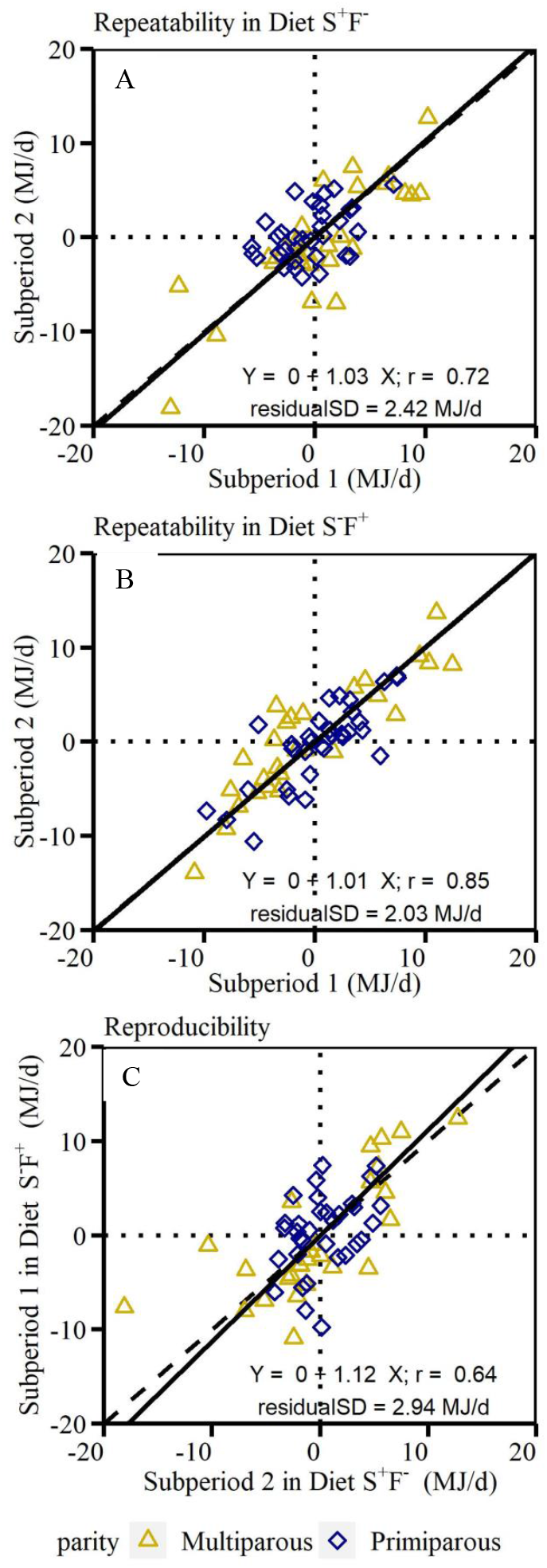
Relationship between feed efficiency estimated within the same diet for repeatability estimation within diet S^+^F^-^ (A), S^-^F^+^ (B) or across diets for reproducibility estimation (C). The dashed black line stands for the first bisector, and the solid black line stands for the least rectangles regression.

**Table 3.** Error of repeatability and reproducibility, and Lin’s concordance correlation coefficient within diet for repeatability and across diets for reproducibility (Repro.) for feed efficiency.

|  | Repeatability<sup>1</sup> |  | Repro.<sup>2</sup> | P-value |  |  |
| --- | --- | --- | --- | --- | --- | --- |
|  | Diet S<sup>+</sup>F<sup>-</sup> | Diet S<sup>-</sup>F<sup>+</sup> |  | S<sup>-</sup>F<sup>+</sup> vs S<sup>+</sup>F<sup>-</sup> | S<sup>-</sup>F<sup>+</sup> vs Repro. | S<sup>+</sup>F<sup>-</sup> vs Repro. |
| SD (MJ/d) | 2.401 | 2.014 | 2.953 | 0.06 | <0.01 | 0.02 |
| CCC | 0.72 | 0.85 | 0.64 | 0.19 | 0.05 | 0.55 |

All cows were first fed with the S<sup>+</sup>F<sup>-</sup> diet (S<sup>+</sup>F<sup>-</sup>: high in starch and low in fibre) and then with the S<sup>-</sup>F<sup>+</sup> diet (S<sup>-</sup>F<sup>+</sup>: low in starch and high in fibre). CCC: Lin's coefficient of correlation of concordance; SD: standard deviation of the residuals of the analysis of variance defined in model 3. <sup>1</sup>Repeatability was estimated within diet using two repetitions of 36 days within diet. Repeatability was defined as the standard deviation of the residuals of model 3. The lower the error of repeatability, the more repeatable it is. <sup>2</sup>Reproducibility was estimated using two repetitions of 36 days (1 repetition/diet): the last 36 days of diet S<sup>+</sup>F<sup>-</sup> and the first 36 days of diet S<sup>-</sup>F<sup>+</sup>. Reproducibility was defined as the standard deviation of the residuals of model 3. The lower the error of reproducibility, the more reproducible it is.

### Effect on feed selection and feed efficiency

The change in diet composition was associated with a change in cows’ feed sorting behaviour. Indeed, the first plan of the PCA (Figure 3) showed that the individual refusals were clustered around the origin of this first plan, close to the two diets samples, and slightly shifted to the more fibrous feed ingredients of the diets (leaves part of maize silage, and straw). This suggests that the cows may have left more fibrous ingredients than concentrates or grains in the refusals, and that individual refusals seem to be different to any particular feed ingredient in the diet. Despite the high number of spectra length waves used to describe the spectrum of each sample, the two first principal components of the PCA explained 94% of the total variability of the spectra of the refusals and the feed samples (Figure 3). When focusing only on the refusals samples (second PCA), the first plan explained 90% of the total variability of the refusals’ spectra (Figure 4). This focus on refusals’ spectra only showed that the refusals were clustered in 2 groups identifiable as the 2 diets (Figure 4). This suggests that the refusals reflect the composition difference between both diets. Within each diet, the refusals spectra differed across cows, but this difference was not associated with feed efficiency differences. Indeed, the randomRNEI was evenly distributed within diet with no clear clustering within diet that was associated with randomRNEI. Moreover, the principal components were poorly correlated with randomRNEI with a correlation of - 0.05 between randomRNEI and principal component 1, and of 0.009 between randomRNEI and principal component 2. If some variability exists in the composition of refusals due to feed sorting, this selection was not associated to feed efficiency differences in this trial neither when cows were fed with the S^+^F^-^ diet nor when fed with the S^-^F^+^ diet.

**Figure 3.**
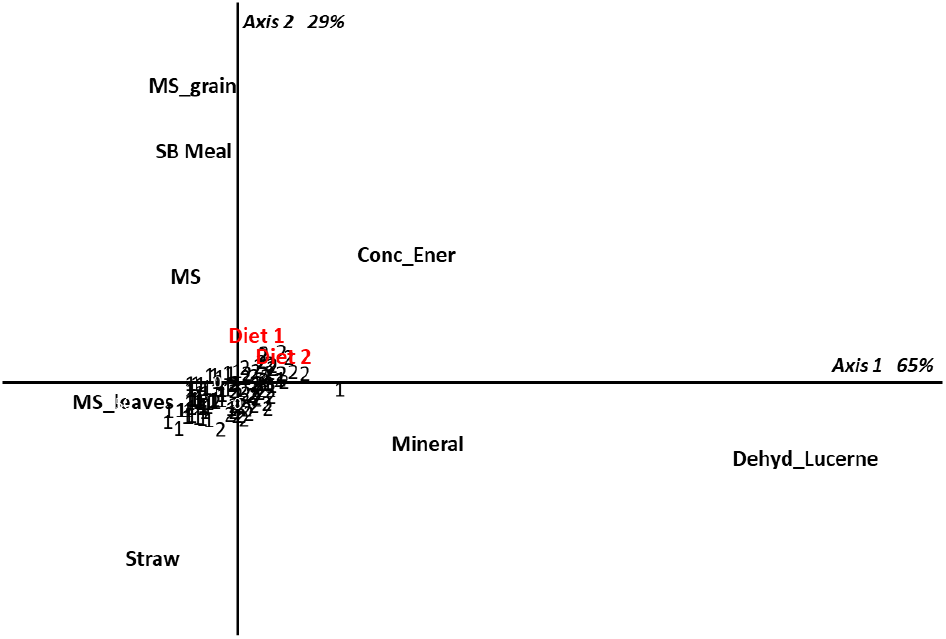
Principal Component Analysis of NIR spectra of the feed ingredients (MS=Maize Silage, MS_leaves = leaves and cane of Maize Silage, MS_grain=Grain of maize silage, Conc_Ener=concentrate energy, SB Meal = soybean meal, Dehyd_Lucerne = dehydrated Lucerne, Mineral), diets (Diet 1 = diet S^+^F^-^ and Diet 2 = diet S^-^F^+^) and refusals of diet S^+^F^-^ (1) and diet S^-^F^+^ (2). The 2 first components explain 94% of the total variance. Diet S^+^F^-^: diet high in starch and low in fibre; Diet S^-^F^+^: diet low in starch and high in fibre.

**Figure 4.**
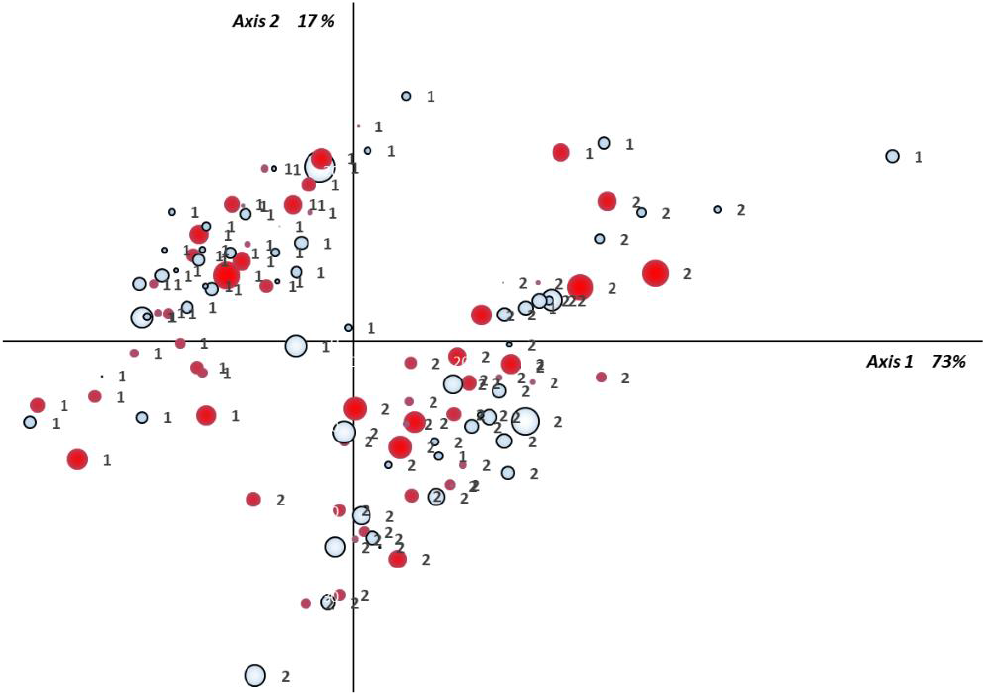
Principal Component Analysis of NIR spectra of the refusals of diet S^+^F^-^ (1) and diet S^-^F^+^ (2). The 2 first components explain 90% of the total variance. The size of the points is proportional to the absolute value of the residual net energy intake and the color is red if randomRNEI is positive (inefficient cows) and blue if randomRNEI is negative (efficient cows). Diet S^+^F^-^: diet high in starch and low in fibre; Diet S^-^F^+^: diet low in starch and high in fibre.

## DISCUSSION

### Feed Efficiency was less reproducible across diets than within diet

Feed efficiency was less reproducible across diets than within diet when using the method defined by ICAR (JCGM, 2012) based on the estimation of errors. However, as shown with the CCC and the cow’s ranking (see section “Availability of data and material” for this supplementary material), the change in cow’s ranking was similar when comparing between the two diets than when comparing within diet over subsequent lactation stages. This shows that the change in diet affected the absolute value of feed efficiency but not cow’s ranking.

This difference between the results observed with the error method and those observed with the CCC shows that it is therefore important to use several indicators when the objective is to characterize reproducibility and repeatability of a method, here of feed efficiency. In our study, we showed that the error of reproducibility across diets was significantly higher than the error of repeatability within diet, while the CCC were similar when comparing across diet and within diet S^+^F^-^, but different when comparing across diets and within diet S^-^F^+^. With one indicator we would have concluded that diet has a significant effect on feed efficiency repeatability, while with the second we would have concluded that the effect of diet does not seem significant. The two indicators are complimentary: the CCC will give the strength of the relationship between the two diets feed efficiency without any information about the dispersion of the residuals of this relationship, while the errors give the dispersion of the relationship. Most of the studies characterizing the reproducibility of feed efficiency in cattle used coefficient of correlations and percentages of cows which changed their efficiency group (Durunna et al., 2011; Potts et al., 2015; Asher et al., 2018). Conclusions based on animals’ re-ranking are highly subjected to the definition of efficiency groups, which is variable across studies. The conclusions about repeatability may even differ with different definitions of efficiency group. We therefore preferred not to use re-ranking to characterize reproducibility of feed efficiency.

Upon the objective of cow’s selection based on feed efficiency, one will prefer one indicator or the other. For example for selection purposes, one may especially be interested in the rate of cows able to maintain their efficiency class, and therefore use the CCC indicator. With this in mind, our results and the literature show (Durunna et al., 2011; Potts et al., 2015; Asher et al., 2018) that the ranking is quite similar within diet over time and across diets, and therefore that the risk to misidentify the most or least efficient cows is minimal. If the objective is to improve efficiency on an existing herd, one may prefer to use the errors indicator. With this in mind, our results show that a diet change affects the efficiency of the herd because the errors are significantly higher when predicting feed efficiency after a diet change than within the same diet. Indeed if efficiency would be reproducible when changing diet’s composition, all cows should behave the same way and their efficiency would be easily predictable with their previous efficiency. This is not observed, because when randomRNEI in first sub-period within diet S^-^F^+^ was predicted with the randomRNEI in second sub-period within diet S^+^F^-^, the regression was able to explain 41 % of the total variability of randomRNEI in first sub-period within diet S^-^F^+^. This low coefficient of determination is mostly explained by the diversity of adaptation of each cow’s DMI to the diet change (Figure 2). Indeed, for similar randomRNEI and similar DMI there were cows which decreased their DMI and decreased their randomRNEI, while others increased both their DMI and randomRNEI, and others maintained both their DMI and randomRNEI. This shows that cows having similar feed efficiency and intake on a specific diet, will not necessarily end up with similar efficiency and intake on a different diet.

To conclude about reproducibility, one should also estimate repeatability and compare it to reproducibility. If the reproducibility is similar to repeatability, the method or estimation is highly reproducible. If the reproducibility is significantly worse than repeatability, the method is less reproducible than repeatable, and therefore the method or estimation is sensitive to the environment. The estimation of repeatability is therefore essential when characterizing the reproducibility of a method or estimation. The lack of confidence interval or statistical test to compare the reproducibility and repeatability indicators in most of the studies (Durunna et al., 2011; Potts et al., 2015; Asher et al., 2018) limits the possibility to conclude about the reproducibility of feed efficiency.

### Similar reproducibility results than in literature

The CCC comparing efficiencies before and after diet change was significantly lower when compared with the CCC of repeatability within diet S^-^F^+^ for feed efficiency (p = 0.05), but was not different when compared with diet S^+^F^-^ CCC (p > 0.1). The lack of significance observed when comparing the reproducibility CCC when diet changed with the repeatability CCC estimated within diet S^+^F^-^ can be explained by the lower repeatability of feed efficiency observed when cows were fed with diet S^+^F^-^. The lower repeatability of randomRNEI within diet S^+^F^-^ can be explained by the lower repeatability observed for NEI within diet S^+^F^-^ as its error of repeatability was 4.60 MJ/d within diet S^-^F^+^ and 5.82 MJ/d within diet S^+^F^-^. The observed decrease in correlation for feed efficiency under reproducibility conditions was also found in literature with Pearson’s correlations of 0.54-0.70 in heifers and 0.42 in steers within diet, that were larger than those observed after diet changed with correlations of 0.40 for heifers and 0.33 for steers (Durunna et al., 2011; Cassady et al., 2016). In dairy cows, correlations averaged 0.65 within diet and 0.56 across diets (Potts et al., 2015). The correlations they observed were lower and closer together compared to the correlations observed in the current study. The higher correlations observed in our study can be explained by the longer period used, that is 36 days, compared to Potts et al. (2015) who used 7 days per sub-period to estimate repeatability correlations within diet. In fact, the longer and the closer to middle of lactation the period was, the better the RFI approximates a RFI estimated over the full lactation (Connor et al., 2019). It would be worth to look at the period length required to reach the maximum repeatability within diet and within lactation for RFI, and then apply this length within diet and estimate reproducibility of feed efficiency when changing diet.

### Diet and Period Effects on Performance

Dietary starch and fibre concentration modified intake and performance in lactating dairy cows. Diets high in starch and lower in fibre compared to diets low in starch and higher in fibre do not necessarily increase intake (Boerman et al., 2015; Potts et al., 2015; Karlsson et al., 2018). Similarly, in the current study, intake, expressed as DM, was not significantly higher when cows were fed with S^+^F^-^ diet than when they were fed with S^-^F^+^ diet (p = 0.26). However, when expressed as net energy, cows ate more when fed with the S^+^F^-^ diet than when fed with the S^-^F^+^ diet (p < 0.01), such as observed in Karlsson et al. (2018). Decrease in dietary starch and fibre concentrations reduced milk production (p < 0.01), as observed in literature (Boerman et al., 2015; Potts et al., 2015; Karlsson et al., 2018), but in the current study this effect was confounded with the effect of lactation stage. This decrease in milk production subsequent to a decrease of dietary starch and increase of dietary fibre is usually observed with an increased milk fat concentration (Boerman et al., 2015; Potts et al., 2015; Karlsson et al., 2018). This was not observed in the current study because neither milk fat nor milk protein concentrations were different between diet S^+^F^-^ and diet S^-^F^+^ (p = 0.56 for milk fat and 0.88 for milk protein). One could argue that our change in NDF between both diets, of 16%, was too low compared to the 18-38% across Boerman et al. (2015), Potts et al. (2015) and Karlsson et al. (2018) to see significant changes in milk fat concentrations. However, in Boerman et al. (2015) only the lowest increase in NDF (18%) had a significant change in milk fat concentration; the other diets having a higher change in NDF did not significantly increase the milk fat concentration. Therefore the increase in NDF may not systematically increase milk fat concentration. This steadiness of milk solids concentrations, combined with the decrease in milk production, resulted in lower net energy exported in milk (p < 0.01) and lower milk fat and protein productions (p < 0.01) on diet S^-^F^+^. Having a diet higher in net energy concentration should have led to more body reserves replenishment and less body reserves mobilization, as observed in Boerman et al. (2015) and Potts et al. (2015). In the current study, a higher dietary starch concentration was associated with lower BW loss (p < 0.01), and similar BW gain (p = 0.28). Those conclusions differences for change in milk solids and BW gain between our study and literature results can be explained by the difference in experimental design. Indeed, the current study was based on a sequential design while the other studies were based on a crossover design (Boerman et al., 2015; Potts et al., 2015; Karlsson et al., 2018). The main limit of a sequential design, as chosen in the current study, is the confounding between the effect of stage of lactation and the effect of treatment, which does not exist in a crossover design. However, crossover designs have limits too: it is impossible to characterize the effect of interaction between treatment and time or to characterize a possible remnant effect of the first treatment on the following treatments. It is therefore difficult to conclude if diet change or just lactation stage explained the decrease in NEI and in milk production in the current study. However, given that cows should replenish their body reserves more and mobilize them less with increasing lactation stages, we can hypothesize that the lack of change in BW gain and the increased BW loss is mostly due to decreased dietary starch concentration. The observed increase in BW and BCS in the current study may be explained by more advanced lactation stages, and less by diet change. Reducing starch and increasing fibre concentrations of the diet increased methane yield, as per kg DMI (p = 0.03) and as per kg milk produced (p < 0.01). This increase of methane per kg milk produced is partly explained by a lower milk production with similar feed intake. The increase of methane yield per kg of DM is more associated with the diet composition. By decreasing starch concentration in the diet and increasing dietary fibre concentration with increased wheat straw and lucerne, the production of volatile fatty acids in the rumen may have shifted in favour of acetate or butyrate profile instead of propionate profile. This shift is usually associated with a friendlier methanogenic environment (Knapp et al., 2014). The expected higher methane yield with lower dietary starch concentration, as per kg DMI or per kg milk, as observed in the current study, has also been observed in literature (Bougouin et al., 2018) but was not consensual. For some research, methane yield per kg DMI did not differ when changing dietary starch concentration (Hatew et al., 2015; Pirondini et al., 2015). A decrease in dietary starch concentration has usually been associated with an increase in methane emitted per day (Pirondini et al., 2015; Bougouin et al., 2018), which was not significant in the current study (p = 0.41). These diverging results about effect of dietary starch concentration on methane production may reflect differences in the effect of dietary starch concentration on DMI, milk yield and diet digestibility. Higher methane emissions per day were observed when the change in dietary starch concentration was associated with significant differences in DMI (Bougouin et al., 2018) or significant differences in diet digestibility (Pirondini et al., 2015; Bougouin et al., 2018). The significant change in methane yield per kg DMI was observed when the change in dietary starch concentration was associated with a change in DMI and a change in diet digestibility (Bougouin et al., 2018). The conclusions on dietary starch concentration on methane emissions seem to depend on the effect of diet composition on both, DMI and diet digestibility.

### Limits of the study

Given that digestibility partly explains feed efficiency differences in lactating dairy cows (Oliveira et al., 2016; Potts et al., 2017), it would be interesting to evaluate the cows’ ability to maintain their efficiency for diets with decreased DM digestibility or for diets with greater physically effective fibre affecting rumen fill. For example, a future study could compare a classic highly digestible fibre, as S^+^F^-^ diet in current study, with a diet including fresh grass or hay.

In the current study, the effect of diet was confounded with the effect of lactation stage because all cows went through the same sequence at the same time. The change in performance, as well as in intake, could therefore be attributed to the diet, but also to lactation stage. This experimental design can also lead to remnant effects of the previous treatment to subsequent periods: results could have been different if diet sequence would have been reversed between the two periods. Repeatability errors in diet S^+^F^-^ were similar to repeatability errors in diet S^-^F^+^ (p = 0.06). Feed efficiency was therefore as repeatable in diet S^+^F^-^ as in diet S^-^F^+^ (Table 3), which suggests that there was no remnant effect of diet S^+^F^-^ on feed efficiency under diet S^-^F^+^ and that lactation stage did not affect feed efficiency variability in our study. Another way to tackle confusion between lactation stage and treatment would have been to adopt a crossover design. However, this design can possibly lead to an interaction between time and treatment, which is not quantifiable, or leads to a remnant effect of the first treatment. Moreover, a similar study was conducted by Fischer et al. where a cross-over design (paper under review) was used. The results and conclusions were similar to the current study. This supports the validity of the current paper.

A last limit to the study is the length of the adaptation period between diet S^+^F^-^ and S^-^ F^+^. When comparing sub-period 2 of diet S^+^F^-^ with sub-period 1 of diet S^-^F^+^, we compared two periods which had different length of adaptation period. Indeed the first had at least sub-period 1 of diet S^+^F^-^ (36 days) whereas the second had 23 days. As we commonly consider that 2 to 3 weeks are enough to ensure that the cows are fully adapted to a new diet, we considered that the adaptation to the diets was achieved for the data used in the current study, and therefore we considered that the difference in length of the adaptation period did not influence the results.

## CONCLUSIONS

The study estimated the reproducibility of feed efficiency in dairy cows when changing the diet concentration in starch and fibre, by comparing the reproducibility across diets with its repeatability within diet. The results showed that feed efficiency was significantly less reproducible when changing diet’s starch and fibre concentration than repeatable within diet over subsequent lactation stage. However the change in feed efficiency ranking of dairy cows was not significantly different when comparing the ranking when cows were fed with the two different diets than when comparing the ranking when cows were fed with the same diet over subsequent lactation stages. The diet change in starch and fibre concentration did not affect the ranking of dairy cows more than does the lactation stage. The change in diet composition affected the feed sorting behaviour of the dairy cows. However this feed sorting was neither related to feed efficiency differences nor to the change in feed efficiency subsequent to diet change.

## LIST OF ABBREVIATIONS

BCS: body condition score;
BW: body weight;
CCC: Lin’s coefficient of correlation of concordance;
CH4: methane;
DM: dry matter;
DMI: dry matter intake;
S^+^F^-^: diet high in starch and low in fibre;
S^-^F^+^: diet low in starch and high in fibre;
NEI: net energy intake;
randomRNEI: random residual net energy intake;
SD: standard deviation.

## DECLARATIONS

### Ethics approval

The protocol has been approved by the ethical committee and the French Ministry of Higher Education, Research and Innovation (Authorization of the French Ministry of Higher Education, Research and Innovation reference APAFIS 3122-2015112718172611).

### Consent for publication

Not applicable.

### Data, script and code availability

The datasets generated and/or analysed during the current study (“dataset_origin.tab”, “dataset_nirs_refusals_v2.tab”), the SAS code (SASscript.txt”), the R scripts (“Rscript_datasetcreation_analysis.R”and “Script_PCA_refusals.R”), and the Metadata (“description_variables_depositpaper.tab”) are available in the following data.INRAE repository: https://doi.org/10.15454/FHRTWJ. In this repository you will also find the supplemental table with the cows’ ranking per sub-period (see table “RFI.tab”).

### Author’s contributions

PF designed the experiment and monitored it; AF gathered and cleaned the data; PG contributed to define the methodology used to check the hypothesis of feed sorting, to analyse and interpret NIRS analysis of feeds and refusals; AF and PF analysed the data and interpreted the results; AF was the major contributor in writing the manuscript; AF and PF read and approved the final manuscript.

## Acknowledgements

The authors warmly thank technical staff and managers at the INRA Méjusseaume research facility who helped managing this experimentation, did a great job in doing precision monitoring of the data, extracting and preparing the data and for being very helpful for biologically clarifying abnormal data.

Version 3 of this preprint has been peer-reviewed and recommended by Peer Community In Animal Science (https://doi.org/10.24072/pci.animsci.100016).

